# Fused regression for multi-source gene regulatory network inference

**DOI:** 10.1101/049775

**Authors:** Kari Y. Lam, Zachary M. Westrick, Christian L. Müller, Lionel Christiaen, Richard Bonneau

## Abstract

Understanding gene regulatory networks is critical to understanding cellular differentiation and response to external stimuli. Methods for global network inference have been developed and applied to a variety of species. Most approaches consider the problem of network inference independently in each species, despite evidence that gene regulation can be conserved even in distantly related species. Further, network inference is often confined to single data-types (single platforms) and single cell types. We introduce a method for multi-source network inference that allows simultaneous estimation of gene regulatory networks in multiple species or biological processes through the introduction of priors based on known gene relationships such as orthology incorporated using fused regression. This approach improves network inference performance even when orthology mapping and conservation are incomplete. We refine this method by presenting an algorithm that extracts the true conserved subnetwork from a larger set of potentially conserved interactions and demonstrate the utility of our method in cross species network inference. Last, we demonstrate our method’s utility in learning from data collected on different experimental platforms.

## 1 Introduction

As the volume and variety of genome scale data continues to increase in quantity and quality, the goal of accurately modeling gene regulatory networks has become attainable [5, 9, 7]. Large-scale data collection efforts have contributed to the development of high quality networks which accurately recapitulate biological processes, but most processes and organisms remain uncharacterized at the network level. Furthermore, as new technologies are developed and some old ones are replaced, such as RNAseq and microarray, it becomes important to be able to combine data from multiple platforms, lest we lose valuable information from existing studies. The problem of inferring related – but not necessarily identical – structure from related – but not identical – data is ubiquitous in biology. Multi-source network inference has applications for learning multiple networks in related species, for learning networks associated with distinct processes within the same species, and for learning networks based on heterogenous data sources. Moreover, as it becomes possible to learn genome-wide regulatory networks, we can begin to compare and to test whether there is conservation of networks across species and biological processes. Our use of model organisms to study biological processes and diseases relevant to humans relies on the assumption of conservation; yet this has not been effectively tested at the genome scale.

We present two methods for network inference based on linear estimates of gene expression dynamics, extending existing dynamical-systems methods for network inference [5, 1, 60]. The core of both methods is the observation that biological information about the relatedness of genes can be used to select which network coefficients should be similar to one another in a multi-source network inference problem (ie orthologous TFs should regulate orthologous genes), and that these constraints can be efficiently represented as penalties in a least-squares regression problem. Taking into account the similarity of putatively conserved interactions improves our ability to accurately describe TF-gene relationships on a genome-wide scale.

Our first method – fused ridge – uses an L2 penalty on the differences between a priori similar interactions (termed fusion penalty), and is useful where the relationships between networks (similarity of genes between data sets) is reliable. In the case where both networks contain an identical set of genes and TFs, this approach can be thought of as parametrically interpolating between treating the data sources separately and combining them together. In the case of multi-species, simply combining two datasets is both unwise – because the networks may differ substantially – but also potentially impossible, because the set of common of genes may be small. Our method allows useful pooling of data even when the overlap between genes is incomplete, or when orthology assignments depart from a strict one-to-one mapping. Our second method – adaptive fusion – uses a non-concave saturating fusion penalty to simultaneously infer the constrained networks and to learn which constraints should be relaxed (ie which parts of the network are genuinely different). With this approach, we seek to identify both conserved and divergent interactions between related networks.

In the case of multiple species, numerous studies have shown that functional conservation exists in gene regulatory networks even across large evolutionary distance [51, 21, 56, 12]. In our fused L2 approach, we assume that for closely related species, orthologous TFs could exert similar regulatory effects on orthologous target genes. These orthology relationships form the basis of a set of constraints which favor – but do not require – networks in which orthologous transcription factors regulate orthologous genes. As a result, data in one species can improve network inference performance in another species (and vice versa). This general framework for multi-species network inference can be extended to an arbitrary number of distinct organisms, each contributing data with only partially overlapping sets of genes. This is an advantage over existing approaches to multi-species network inference, which infer only a sub-network for which orthologs exist in every species [26].

This approach of introducing constraints on the similarity between specific regulatory interactions can be extended beyond the case of multi-species network-inference from orthology; any biological prior on similarity of regulation can be used in place of orthology. For example, we can use fused regression to combine datasets obtained using different platforms or experimental techniques and we can introduce constraints that favor genes in the same operon (or having similar promoters) towards having similar regulators.

Existing multi species approaches often use orthology as a proxy for functional conservation [50, 46, 26, 27, 62], or attempt to learn functional similarity via expression data [17]. Orthology can be approximated using readily identifiable sequence similarity, which is often a useful predictor of functional similarity [59, 24]. In multi-species network inference, our fused L2 approach minimizes a cost function that strives to simultaneously fit expression data in each species and produce networks that are consistent with evolutionary constraints created using orthology. However, many genes will have evolved different functions and therefore may have new regulatory interactions. For example, gene duplications may lead to neofunctionalization [11] of the duplicated genes. In the case of comparing networks from related cell lines from the same species, changes in chromatin configuration may affect our hypotheses about the similarity of interactions between pleiotropic TFs and target genes across cell types (a within-species analog to neo-functionalization) [35]. Identifying interactions that are present in one species but not another is of direct biological interest, but existing approaches to network inference are unable to effectively test the hypothesis of conserved subnetworks. Observing a large difference in the weights of regulatory interactions obtained though independent inference of multiple networks is perhaps the best (least biased) evidence against conservation of orthologous regulatory interactions (cases where target and regulator have orthologs across species). However, this is sometimes weak evidence, as network inference is typically under-constrained [39], meaning there could be a different set of networks for which conservation does hold, and which fit the data almost as well. We propose using our adaptive fusion approach to simultaneously perform network inference and evaluate edge-conservation (or lack thereof).

We approach this problem by introducing a saturating penalty function based on statistical efforts to develop unbiased regularization penalties for fused regression [62, 13]. The main difference between the L2 fusion approach and our new adaptive fusion approach occurs when the difference between presumed analogous interactions is large despite the fusion penalty. We assume that cases where the method is unable to reconcile the likelihood and the fusion constraint derived from orthology (in the case of multi-species fusion) or identity (in the case of multi-platform fusion) correspond to cases with evidence of divergent TF-to-target-gene interactions. Practically, this inability to reconcile likelihood and conservation hypothesis manifests as large differences in model weights across species or platforms. We account for this possibility in our adaptive fusion model with a relaxation of the fusion penalty in cases with extreme model weight divergence. The resulting cost function is non-convex and difficult to optimize [13]; however, we can approximate its solution and obtain deeper insight into functional similarity than is available through strict orthology enforcement or the comparison of separately learned networks.

Although the fusion constraints we employ can be described as arising from orthology - which links genes - it is important to note that the constraints themselves link individual regulatory interactions. This finer level of representational granularity is critical to the functioning of adaptive fusion, and means the method can accommodate any form of prior on expected similarity between regulatory interactions, even priors that cannot be decomposed into gene to gene mappings. We develop two algorithms for solving efficiently multi-output least-squares regression problems with pairwise L2 fusion penalties on entries of the coefficient matrix. We also introduce - in the form of adaptive fusion - the idea of a saturating penalty function on fusion constraints, and estimate the solution to the resulting optimization problem through iterative application of the fused L2 algorithm.

We test the ability of fused L2 and adaptive fusion to improve network recovery on both synthetic data and by comparing related networks in the bacteria species *Bacillus subtilis* and *Bacillus anthracis*. This shows the applicability of our method in combining different datasets and leveraging similarity across organisms as well as within a network in order to improve network inference. We explore the circumstances under which each approach is optimal, and evaluate the robustness of adaptive fusion to incorrect orthology, simulating the biologically relevant cases of neo-and sub-functionalizations.

## 2 Methods

### 2.1 Statistical approach and background

We consider prediction and coefficient estimation problems with *N* observations of *M* dependent variables *y*1,1,*y*2,1,…*yN*,1,*yN*,2,…,*yN,M* and *p* features *x*_*i,j*_, *i* = 1,2,…,*N,j* = 1,2,…,*p*. We begin with a standard linear regression model:

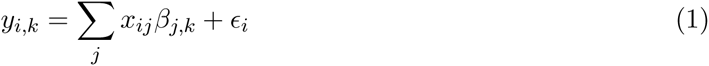

with errors *∊*_*i*_ having mean 0 and constant variance, and predictors *xij* having mean 0 and unit variance. We are interested in the case where *p* > *N*. Many methods have been proposed to deal with the underconstrained case, and have been applied to genomic data [57, 36]. For example, ridge regression penalizes the L2 norm of the coefficients *β*_*1,j*_ in order to avoid overfitting [22], and can be thought of as a mean-zero Gaussian prior on the coefficients. More complicated penalties have been developed to represent specific expected or desireable structure in a regression model’s coefficients. For example, Land and Friedman [32] proposed a fusion penalty which encourages smoothness of the estimated parameter vector. Previous approaches have used fusion penalties to draw statistical strength across multiple regression tasks [29, 33, 8, 47, 20]. Price et al. and Bilgrau et al. use a fused ridge estimator for jointly estimating multiple inverse covariance matrices [49, 4]. We take a related approach to these prior works, adding an L2 penalty on the differences between coefficients to the existing ridge penalty in order to incorporate prior knowledge about relationships between input-output pairs:

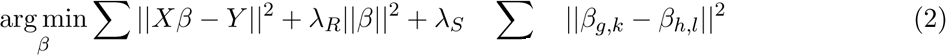

where *X*, *Y*, and *β* are matrices, and *β*_*g,k*_ ≈ *β*_*h,l*_ denotes fusion between entries of *β* (enforcing similarity between model weights across separate data-sets). Note that, like ridge regression, this penalty can be thought of as representing a Gaussian prior on the coefficients *β*. In the case where *β* is a column vector, introducing this penalty is equivalent to assuming that *β* is sampled from a multivariate Gaussian with inverse covariance matrix 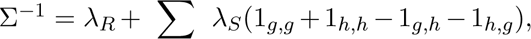
where *I* denotes the identity matrix and 1_*i,j*_ a matrix of zeros with 1 in its *i,jth* entry. In the case of a two-coefficient model with fusion between the coefficients, for example, fused L2 is equivalent to assuming a prior with variance (*λ*_*R*_ + *λ*_*S*_)/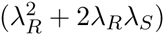 and covariance 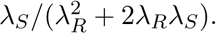

In many cases, however, there is some uncertainty about the relationships that should be enforced. Sohn et al. attempt to simultaneously learn the regression coefficients and the output structure [53]. We develop a similar approach, by applying a penalty function bounded by a constant to produce unbiased estimators for large coefficients, combined with an L2 penalty, similar to SCAD-L2 [61].

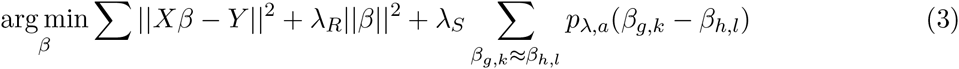

where the penalty *p*_*λ,a*_ has derivative

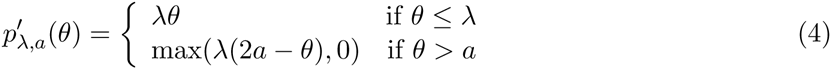

This approach allows us to simultaneously learn the regression coefficients and evaluate the validity of our prior information (this model relaxes the fusion penalty when model components are irreconcilably different).

### 2.2 Application

Although our approach is generalizable to a wide variety of multi-source network inference problems, we begin with the concrete example of network inference in two related species. Our approach to multi-species network inference is based on the hypothesis that gene regulation in related species is governed by similar but not necessarily identical gene regulatory networks, due to conservation of function through evolution. We represent conservation of network function by introducing constraints into the objective function for network inference that penalize differences between the weights of regulatory interactions believed to be conserved. These constraints favor the generation of similar networks for related species, and in the generally under-constrained regime of network inference can improve the accuracy of network recovery. We then go on to introduce a method to test the assumption of conserved network structure, and to relax the associated constraints on pairs of interactions for which the data does not support conservation (where conservation of a regulatory edge is implied by like model weights across data-sets).

### 2.3 Approach overview

#### Algorithm 1 Network inference using fused regression

load expression data

load orthology

create priors and fusion constraints

partition gold standard into training and leave-out

generate TFA matrices using gold standard training set

set *a* if using adaptive fusion

**for** *k* in folds **do**

partition expression data into training and leave-out set

*λ*_*R*_ parameter selection using training set

*λ*_*S*_ parameter selection using training set

run fused regression

return PRC and ROC curves using leaveout gold standard

**end for**

average PRC / ROC curves over folds

### 2.4 Gene regulatory network

We model the transcription rate of each gene as a weighted sum of transcription factor expression, and seek to identify the identities and regulatory weights of these TFs. This formulation matches that of the existing *Inferelator* algorithm, which models gene expression with linear differential equations [6]. Our primary data for learning gene regulatory networks is expression data, consisting of time-series and steady state experiments. The rate at which *x*_*i*_, the observed mRNA expression of gene *i*, changes, is governed by degradation of existing transcripts with rate *α* plus a linear combination of transcription factor (TF) expressions.

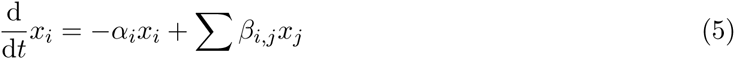

where*β*_*ij*_ represents the weight of TF *j* on gene *i*, and *α* is the decay rate of gene *i*. We fix the decay rate *α* for all genes, and set it assuming a time-constant of 10 minutes [19, 52], as in [18]. Let *x*_*i*_(*t*) be the expression of gene *i* at time *t*. Given time-series data on the expression of gene *i* at timepoints *t*_*k*_ and *t*_*k+1*_, we can approximate the rate of change of *x*_*i*_ as 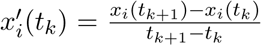. We treat steady-state data as having a derivative of zero. This gives us, for each gene *i* and time *t*_*k*_ an equation

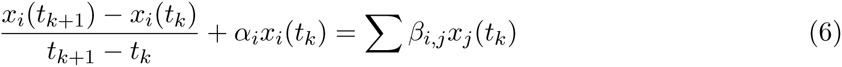

where *j* ≠ *i* for time series and

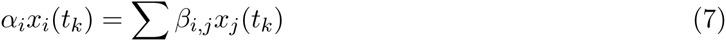

for steady state.

We can summarize these equations in matrix form as

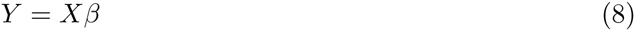

where *Y* is the gene expression matrix, *X* is the TF expression matrix, and *β* is the regulatory weights we are interested in learning. We are interested in learning *β*, the matrix representation of the gene regulatory network, where the weight in a given position represents the regulatory weight of a TF on a gene. Positive weights represent activation, negative weights represent repression, and 0 weights represent the absence of an interaction. The matrix *β* can be solved using linear regression. Because there are typically far fewer conditions than possible regressors (TFs), we introduce a ridge regularization constraint with weight *λ*_*R*_ and solve

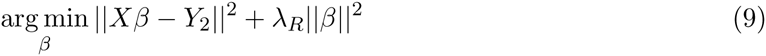

This is similar to the formulation used in the *Inferelator* algorithm, which we extend to the case of simultaneously inferring multiple networks.

Transcription factor expression is not always the best predictor of its gene targets’ expression, so previous network inference methods attempt to estimate transcription factor activities prior to network inference. When there exists a set of prior known interactions, we are able to estimate transcription factor activity (TFA) using network component analysis [37], as in [1, 15], and use TFA as explanatory variables instead of transcription factor expression.

### 2.5 Fused gene regulatory networks

Information about the partially conserved structure of gene regulation is introduced through the incorporation of constraints into the above regression formulation. These constraints penalize differences between interaction weights in the networks of multiple species that are expected to be similar based on prior biological knowledge. We can then solve the penalized regression problems simultaneously, in order to obtain a gene regulatory network (GRN) for each species. Consider the case of organisms *A* and *B*, governed by GRNs *β*^*A*^ and *β*^*B*^ (the following approach applies equally well to more than two species but for simplicity we continue with the case of two species). When TF *g*^*A*^ in organism *A* and TF *h*^*B*^ in organism *B* are orthologs, and gene *k*^*A*^ and *l*^*B*^ are orthologs, then we expect that the *g*^*A*^ ⊒ *k*^*A*^ interaction weight should be similar to the *h*^*B*^ ⊒ *l*^*B*^ interaction weight, and we introduce a fusion constraint between these analogous interactions. In terms of the above regression formulation, we expect that 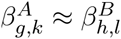, and include a penalty term 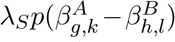 in the quantity being minimized in order to encourage similarity. The function *p*(*x*) controls the shape of the relationship between weight dissimilarity and penalty, while scalar *λ*_*S*_ controls the overall scaling of the penalization of differences between fused coefficients. *λ*_*S*_ controls the tradeoff between fitting the expression data-sets individually and producing a set of networks that conform to evolutionary prior knowledge. This gives us the final equation to be minimized:

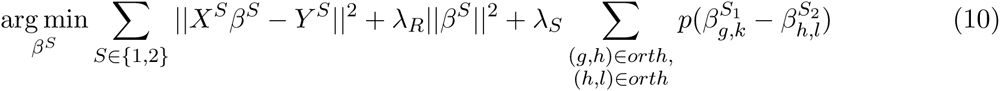

where the second sum is over pairs of interactions with fusion constraints. In the fused L2 algorithm presented here, the penalty function is equal to the L2 norm of the difference in regulatory weight of fused coefficients, *p*(*x*) = *x*^2^. Every component of the objective function is an L2 norm and thus the problem is convex and can in fact be solved through linear regression with an augmented design matrix.

As an example, consider the case where there is a one-to-one orthology between the species being considered (ie different cell-lines of the same organism). The choice of *λ*_*S*_allows one to interpolate between fitting each network independently (*λ*_*S*_ = 0) and pooling data together as if it came from one source (*λ*_*S*_ = inf). In addition to performing well between these extremes, our method allows pooling of data even when there is incomplete orthology. By introducing constraints on the similarity of individual interactions, rather than on the networks as a whole [38], we can pool some information across species even when a small fraction of genes have orthologs.

### 2.6 Adaptive fusion

Fusion constraints penalize dissimilarity between interactions thought to be analogous based on *a priori* knowledge. For example, orthology can be used to predict which interactions will be similar across species. With an L2 fusion penalty, interaction weights which differ from each other by a large amount are excessively penalized, which effectively ensures that fused interactions are assigned similar weights. This will be inappropriate for interactions which are identified based on orthology as being analogous, but which are no longer similar due to evolutionary changes. We propose that a saturating penalty that is relaxed once differences in weights grow beyond a certain point (interactions which appear to be very different based on the data are effectively unfused). A related problem has been studied in the context of LASSO regularization, where it was shown by Fan and Li that using a saturating penalty retains many of LASSO’s desireable properties while removing its bias towards model weights of 0 [13]. They further showed that, although the resulting loss-function is nonconvex, good results can be obtained with a local quadratic approximation of gradient descent. Several saturating penalties, such as SCAD [13] and MCP [62], have been discussed in the context of sparse regression. We introduce a modified form of MCP to the problem of penalizing differences between fused coefficients. The principal difference between the penalty we adopt and SCAD/MCP is that both of these penalties are L1 like at the origin, producing sparse solutions. Some network inference approaches use L1 penalties to produce sparse networks, on the basis that biological networks are thought to be sparse. However, as we are penalizing differences in interaction weights, rather than the weights themselves, there’s no reason to assume that most differences will be exactly zero, and an L2 penalty - equivalent to an assumption that the differences between fused coefficients are Gaussian distributed - may be more appropriate.

We use a penalty on the difference between fused coefficients *θ* which is L2 like at the origin and saturates at *θ* = *a*. Written in terms of its derivative, the penalty *p*^’^_*λ,a*_

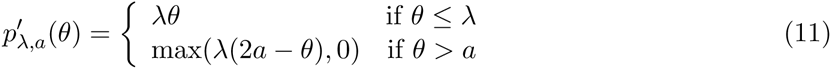

As in [13], we solve using iterative local quadratic approximation. Specifically, *β*^*S*^ (*t*) is the network on iteration *t*. For each fused 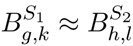 we define:

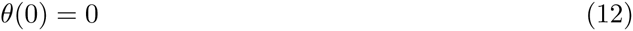

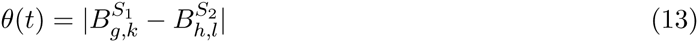

and introduce a fusion constraint 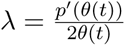

*β*^*S*^ (*t* + 1) is obtained by fitting the ridge-fused model with fusion constraints given by the above *λ*_*S*_. This is useful because all our penalties can be treated as L2 and therefore retain the properties of ridge regression, and can be solved using the fused L2 algorithm we develop.

Our adaptive penalty function introduces, in addition to regularization and fusion penalty weights*λ*_*R*_ and *λ*_*S*_, an unknown parameter *a*. We could employ grid search using cross-validation to search for the best parameters, but for many data sets, this can be computationally expensive. Moreover, we are primarily interested in using this saturating penalty as a way of testing the hypothesis that conservation in GRNs can be predicted based off of known similarities between genes. Therefore, we propose a user-defined *a*, where this parameter is set using the distribution of differences between fused weights from independently fit networks. The choice of which value in this distribution to use for *a* represents the working hypothesis for the fraction of fused interactions which should be unfused.

### 2.7 Solving fused L2 problems using augmented matrices

We begin with the problem of constructing a design matrix to map our problem to that of solving a fused L2 regression problem with a single response variable. We then go on to show that, although the vectorized solution involves solving an impractically large system of equations, under typical biological conditions the structure of constraints allow the problem to be broken up into many smaller subproblems. Key to this approach is the observation that ridge constraints can be incorporated into a least-squares regression problem by appending a scaled identity matrix to the design matrix, and a corresponding number of zeros to the response vector. Similarly, a fusion constraint *λ*_*S*_(*β*_*i*_ − *β*_*j*_)^2^ can be incorporated into a least-squares regression problem by appending a row containing 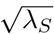 in the ith position, −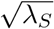 in the *j*th position, and 0s elsewhere to the design matrix, and zero to the response vector. In order to convert an optimization over multiple response variables and multiple sources into an optimization with a single source and response variable, we vectorize as follows: we construct a new design matrix by diagonally concatenating design matrices from relevant regression problems, and create a new response vector by concatenation of corresponding response vectors.

This is equivalent to the original problem due to the block structure of matrix multiplication. In an ordinary regression problem each response variable can be solved independently, and vectorization is unnecessary. However, in fused regression, we append additional rows to the design matrix that link entries of the interaction weight matrix associated with different response variables (figure 8). As a result, these linked response variables must be solved simultaneously through vectorization. Two response variables are linked by a fusion constraint if any of the regulatory weights affecting those genes are linked by a fusion constraint. Two response variables must be solved simultaneously if there is any chain of linked response variables connecting them. However, every other response variable can be solved separately. In biological terms, the regulators of two genes (whether in the same species, or different species) must be solved together if there is a fusion constraint linking those genes’ regulators, or if there is a chain of such constraints. If the networks for a large number of genes are solved simultaneously, the system of equations can quickly become intractable.

**Figure 8:**
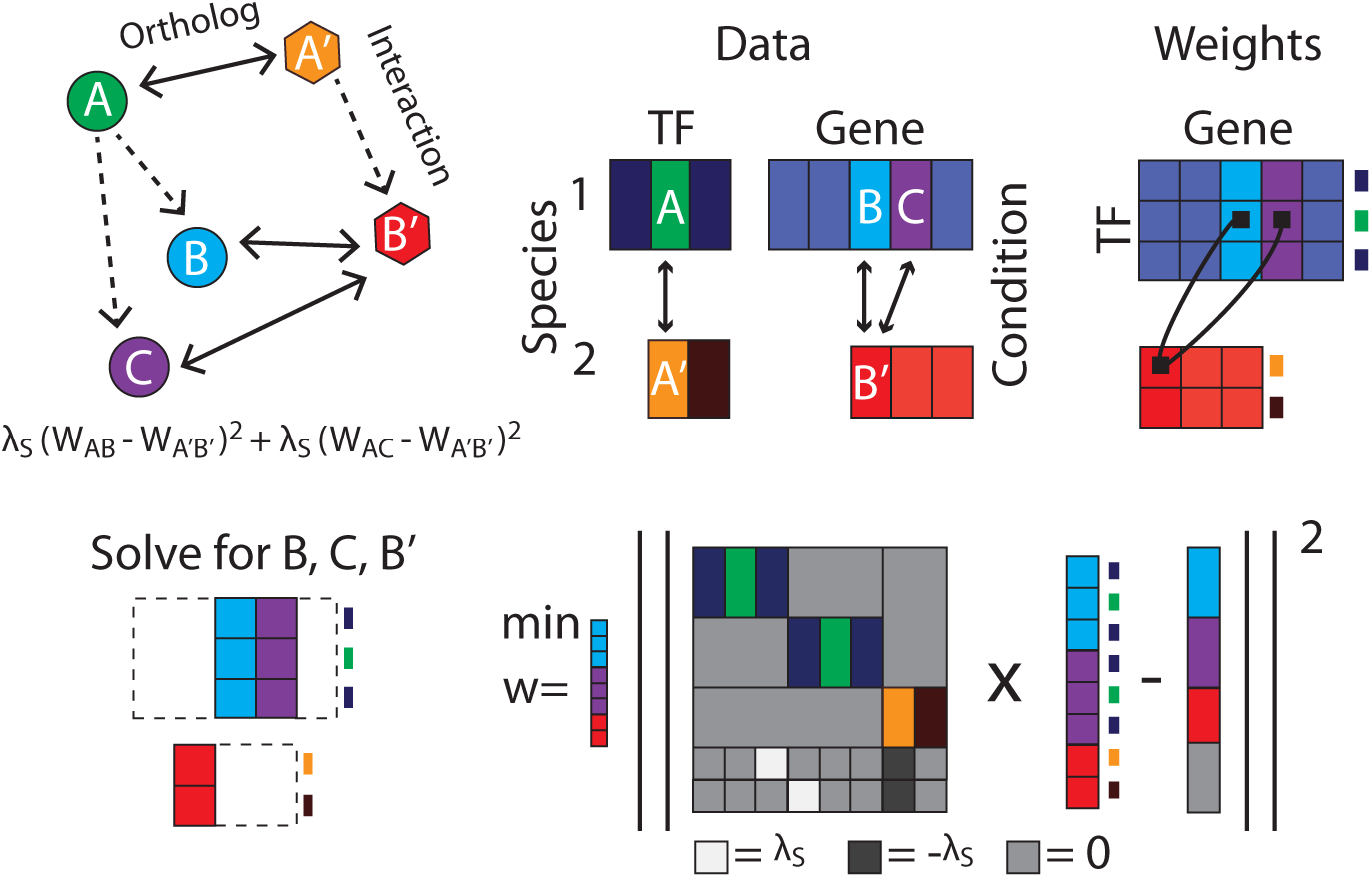
Schematic representation of design matrix construction. Here, the circles and hexagons correspond to different species. Bidirectional arrows represent orthology information and dotted arrows represent putative interactions between TFs and genes. Rectangles represent matrices; because weights can be solved independently unless there exists fusion constraints between them, we identify related weights and construct matrix for solving.

In order to avoid this difficulty, we use depth-first search to identify linked columns of each TF expression matrix, then form design and response matrices through vectorization. We can then incorporate fusion constraints as in the case of single-source single response-variable fused regression. In most cases, we have found the direct solution using augmented matrices to be adequate (possible due to the sparse structure of orthology links; only a small number of genes must be solved at once). In the general case, the size of the design matrix is proportional to the number of response variables that must be solved simultaneously. Because the scaling of this algorithm has a complicated dependence on the constraint structure used, a general description of its runtime is difficult. However, in the case of multi-species network inference with one-to-one orthology, the network associated with each pair of orthologous genes requires solving a linear system with approximately twice as many observations and unknowns as the single species case. Linear systems of this size can be solved quickly using standard techniques, and runtime using our bacterial datasets clocks in around thirty minutes. When the size of the groups of genes linked by fusion constraints becomes large (when organisms have a number of many-to-many orthologous blocks), however, the augmented design matrix approach becomes slower and we discuss further optimisations to this scheme below to enable scaling to these regimes.

### 2.8 Solving fused L2 problems using iterative solver

To address scaling limitations when many-to-many fusion constraint blocks occur, we developed an iterative solver that uses coordinate-wise descent to solve for solutions corresponding to a sequence of values of fusion penalty weights. As our fused L2 method uses a convex and differentiable penalty function, this approach converges to a global minimizer. Although less efficient than the augmented design matrix approach we developed for cases where fusion constraints are primarily one-to-one or few-to-few, the iterative solver has the advantage of computing a solution path for *λ*_*S*_ and scaling well across a wider range of biological applications.

On each iteration *t* the iterative solver computes

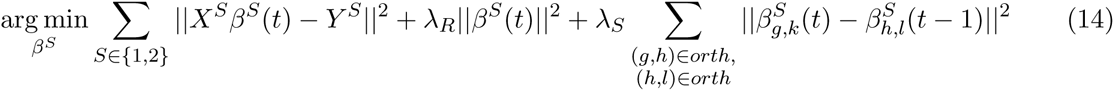

Note that this is almost identical to equation 5, but now the network *β* is a function of the iteration number *t*. On each step, we compute *β*_*S*_ that minimize a penalized cost function where the fusion penalties encourage similarity between a parameter and its fused-to parameter from the previous iteration’s solution. This process is iterated until the estimated *β*s converge. Because each iteration reduces the error between *β*(*t*) and *β*(*t* − 1), and because *β*(*t*) = *β*(*t* − 1) is the globally optimal solution, this process must eventually converge to the same network as equation 2. Although we have not produced bounds on the convergence rate, which also depends on the structure of constraints, in practice a small number of iterations (~10) are necessary.

### 2.9 Fusion and regularization path

Optimizing over both parameters, *λ*_*R*_ and *λ*_*S*_, is computationally prohibitive and we opted to test a heuristic where we optimize the two parameters separately. Our procedure first optimized *λ*_*R*_ with *λ*_*S*_ = 0, then optimized *λ*_*S*_ using this value of *λ*_*R*_. This procedure is guaranteed to achieve the best unfused solution in the case when *λ*_*S*_ is constrained at 0. As a result, any performance gains of fused regression are a lower bound on the highest achievable performance gains. To optimize *λ*_*R*_ we use cyclical coordinate descent algorithms from the ‘glmnet’ package [14] to compute a ridge regularization path. We use cross validation to select the optimal *λ*_*R*_ parameter from this path, selecting the *λ*_*R*_ which minimizes the average error of prediction on a leave out set across cross validation folds. Following selection of *λ*_*R*_, we search for optimal *λ*_*S*_by computing the solution path from the iterative solver (using the sequence of successive model weights) again using cross validation to select the optimal parameter. Note that both parameters are chosen without reference to the gold standard, which is used in a separate evaluation of network quality.

### 2.10 Simulated data

We generate simulated data to evaluate the ability of our fused L2 approach to learn the true network and to show that sharing information between similar but not identical data sources results in more accurate network recovery. Generation of simulated data begins with the production of random orthology mappings with sparcity simlar to that found in real data-sets. We produce a one-to-one orthology by pairing random genes until a specified fraction have been assigned orthologs. This process is carried out separately for TFs and non-TF genes, so that TFs and non-TF genes are never assigned to be orthologous. We then produce a pair of random networks (*β*^1^ and *β*^2^) as follows. For each unfilled entry in *β*^1^ or *β*^2^, we enumerate the set *C* consisting of the entry along with every entry in either matrix to which it is fused. With probability equal to the sparsity rate we assign every entry in *C* to be 0, otherwise we sample a value *ν* ~ *N*(0,1) and independently assign each entry in 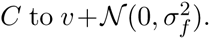 *δ*_*f*_ is a parameter that controls the distribution of differences in the values of fused coefficients, so that the nonzero coefficients of *β*^1^, *β*^2^ are distributed as 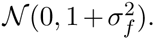

Given a network *β*, we generate *N* samples of gene expressions at two timepoints. The condition by gene expression matrix for timepoint one, *Y*_*T*1_, is sampled randomly from a multivariate Gaussian distribution with identity covariance matrix. *X*_*T*1_ is the TF expression sub-matrix of *Y*_*T*1_, and consists of columns of *Y*_*T*1_ that correspond to TFs. Treating the decay rate as 0, the gene expression matrix at timepoint two, *Y*_*T*2_ is sampled as *Y*_*T*2_ = Y_*T*1_+_*T*1_+∊, where ∊ is a Gaussian noise term. This process is carried out separately for each network. Following generation of simulated data, we may introduce error into the orthology mapping. This can take the form of discarding a specified fraction of true orthologies (governed by a false-negative rate), by introducing random false orthologies (governed by a false-positive rate), or by adding Gaussian noise so that fused interactions are not identical (described above). For convenience, the false-positive rate is specified in units of the number of true orthologs, and not the number of possible orthologs. The list of priors can in a similar fashion be manipulated to include false positives and false negatives.

### 2.11 Ranking regulatory hypotheses

In previous work, betas were rescaled as to form a matrix of confidence scores *S* as follows

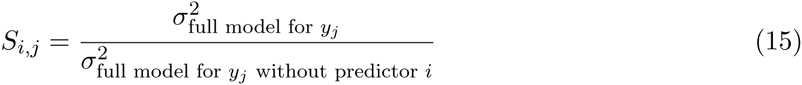

Computing residuals with respect to the data alone would disregard information gained through fusion, because certain interactions may be large due to fusion, rather than their individual explanatory power. Instead, we used an approximation

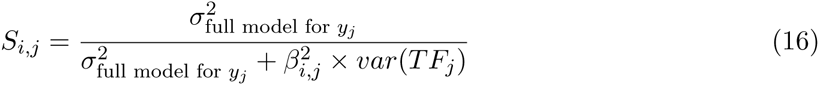

### 2.12 *B. subtilis* and *B. anthracis* data and orthology

We used a dataset collected for PY79, a derivative of strain 168, available on GEO with accession number GSE67023, and a dataset using BSB1, another derivative of strain 168, available at GEO with accession number GSE27219. We used two datasets for *B. anthracis*, transcription profiling during iron starvation (E-MEXP-2272 on ArrayExpress), and time series over the life cycle (E-MEXP-788 on ArayExpress). We ran Inparanoid to obtain orthology mapping for *B. subtilis* and *B. anthracis* [44]

## 3 Results

We used both synthetic networks and real data to test the ability of fused regression to improve the performance of network inference, and the ability of our adaptive fusion procedure to identify conserved interactions between orthologous genes. For the synthetic data, we generated random pairs of networks in which orthologous genes have similar regulatory interactions, and then sampled gene expression from these networks, which we used to derive learned networks for comparison with the input (true) networks. For real data, we computed recovery of a known gold standard in *Bacillus subtilis* [41].

### 3.1 Using fused regression to learn related networks

It is known that the accuracy of network inference improves with additional data [3]. Additional data is not always readily available and does not always make identification of unique network weights possible; but often there is an abundance of related data. Using related data for network inference allows us to leverage statistical power from disparate sources, effectively increasing the sample size and boosting the sensitivity and specificity of learned interactions. We created synthetic networks to approximate two related biological processes, then evaluated performance of our fused L2 regression, which learns the networks simultaneously given a prior on the relatedness of interactions. We compared recovery of the two artificial 10 TFs by 200 genes networks, using fused L2 versus learning networks separately, and varied the amounts of simulated expression data samples made available to the solver. When the amount of data from the second species was held constant, increasing the amount of data available for learning the network for the first species resulted in a more accurate network prediction, as expected (figure 1b). When we increased the amount of data from the second species, we obtained performance gains on network one using fused L2 regression, demonstrating our ability to improve network inference on one dataset through incorporation of a related dataset.

**Figure 1:**
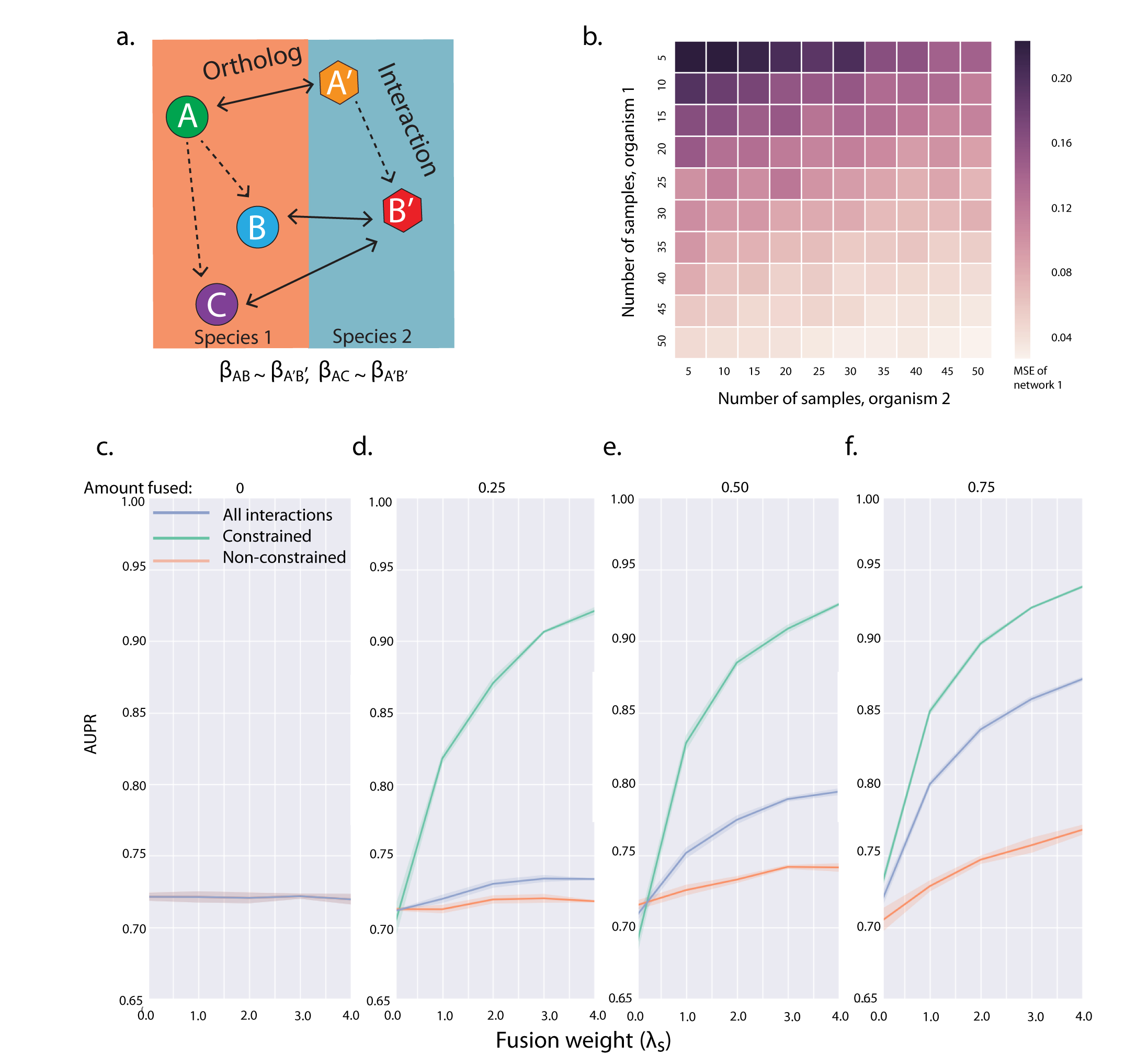
**A**. Schematic representation of the the generation of fusion constraints from orthology mappings. Dashed arrows indicate potential regulatory interactions, while solid arrows denote orthology. We introduce fusion constraints for pairs of interactions for which both the regulator and regulated gene are orthologs of one another. In this example, we would introduce a constraint between the (*A, B*) and (*A*^’^, *B*^’^) interactions and the (*A, C*) and (*A*^’^, *B*^’^) interactions. **B**. In order to demonstrate the utility of fused network inference in combining data, we generate two networks with 10 TFs and 200 genes (75% sparsity). Mean squared error of the inferred vs. true coefficient matrices for network 1 are plotted as a function of the number of conditions generated for species 1 (x-axis) and the number of conditions generated for species 2 (y-axis). As expected, increasing the number of samples available for the species of interest improves network inference performance. However, because we are fusing to data from a related species, similar gains are observed when increasing the amount of data available in this second species. **C-F** Show the varying effects of fusion on simulated networks with different levels of conservation. We generate a series of networks with 20 TFs by 200 genes in two species, (50% sparsity) while varying the fraction of gene orthologies in the simulated networks. For each network, we evaluated AUPR on one of the species for: all interactions (blue line), interactions with fusion constraints (green line), and interactions without fusion constraints (orange line). At every level of conservation, constrained interactions show the largest benefits of fusion, with the magnitude of the benefit growing with fraction of orthologous genes. When the no networks are highly conserved, however, even interactions that are not directly constrained through fusion are recovered more accurately as *λ*_*S*_ increases.

When disparate data is generated from identical processes, the weight assigned to fusion constraints should be very large. On the other extreme, when the networks have diverged considerably, a large fusion weight may impair network recovery. We performed an experiment on simulated data in a multi-species network-inference problem to assess the effect of the similarity of networks, and the effect of the fusion weight on network recovery (figure 2); pairs of synthetic networks with varying similarity were then built. The main factors governing similarity between our generated networks was the extent (and accuracy) of the orthology mapping, and the variability between conserved, interactions (degree of subnetwork conservation). We then conducted a series of simulations to assess the effect of increasing orthology coverage on network recovery. When the conserved subgraphs were very similar - when the interactions in the conserved subnetwork were nearly identical, emulating the case of closely related organisms or similar processes (figure 2a) - increasing the weight of the fusion penalty *λ*_*S*_ improved network recovery. As the size of the conserved subgraph increased, this effect was enhanced. To simulate the case of distantly related organisms, we created networks where the differences between conserved interaction weights were drawn from high variance distribution, mimicking weak conservation (figure 2b). We showed that even for networks where the conserved subgraph was weakly conserved, there exists a model parameterization where fusion regression improves network recovery.

**Figure 2:**
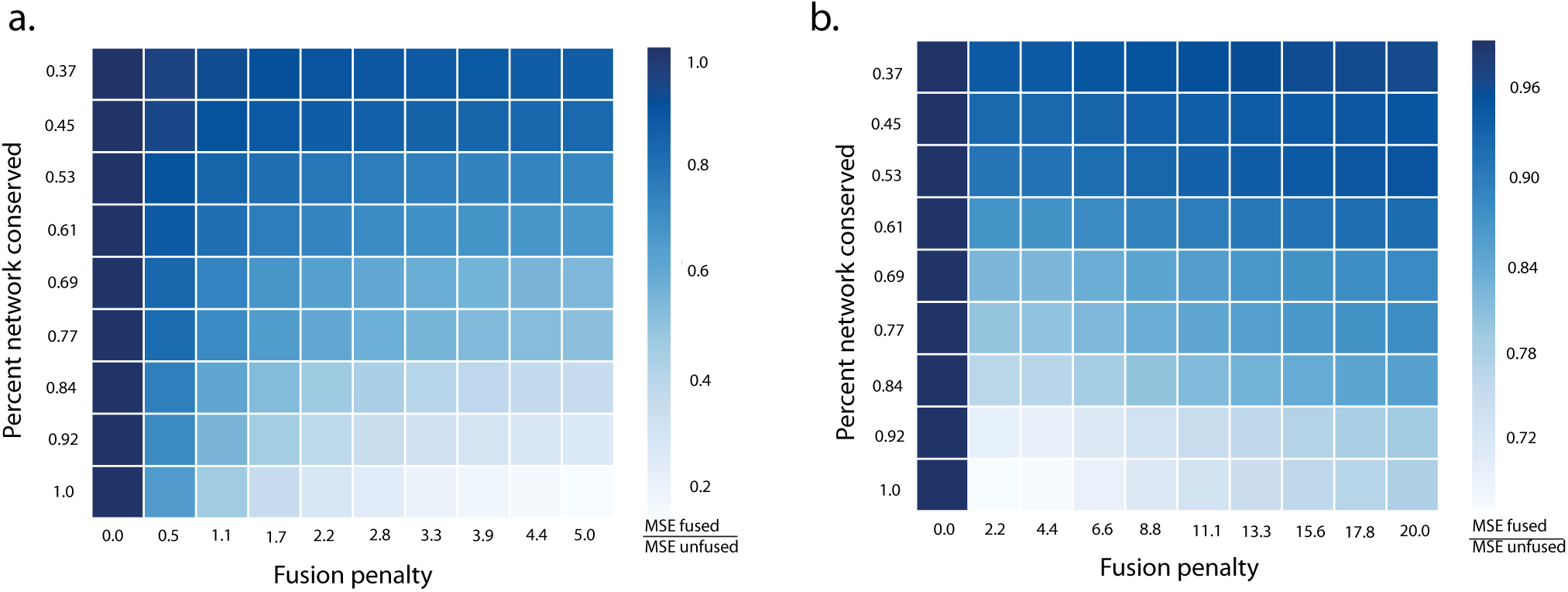
This figure demonstrates the interaction between fusion weight (*λ*_*S*_), the degree of orthology coverage of networks, and the degree of similarity between fused interactions. **A**. We generated a series of pairs of 10 TF by 200 gene networks (75% sparsity) in two species. These networks had minimal fusion noise, so that pairs of interactions linking orthologs had nearly identical weight. We varied the fraction of genes with orthologs (y-axis) and the weight of fusion in network inference (x-axis), and measured performance as the mean-squared error of the true vs inferred network weights. In order to more clearly visualize the varying effect of fusion, performance is plotted in relative units of the of the unfused MSE (left column) for each level of orthology. This was necessary because each row represents a different pair of networks, generated with a different level of orthology, for which baseline performance varies. Performance gains from fusion are, as expected, largest when the degree of orthology is large. However, even when the fraction of genes with orthologs is relatively small (top row), we observe gains from fusion. When fused interactions are nearly identical, larger fusion weights always outperform smaller fusion weights. **B**. Simulates the case where genes which are orthologs may have different regulatory weights. We generated a series of networks as in **A**., but with larger (7.5 ×) gaussian noise added the weights of fused interactions. As in **A**., benefits from fusion were observed at every level of gene orthology. However, unlike **A**., there was an optimal intermediate value of *λ*_*S*_ that traded off between the benefits of fusion and the cost of combining heterogenous data.

### 3.2 Fused regression improves performance on both the constrained and non-constrained parts of the network

Our approach is useful for learning networks from similar sources such as related cell types from the same species, where there exists a one-to-one mapping of genes, as well as datasets where the orthology mapping does not span all genes. This can occur when using different technology, eg microarray and RNAseq, where there is incomplete overlap in the genes that each method assays as well as incomplete overlap in the genes expressed in different experimental designs. When orthology is incomplete we are interested in knowing if performance gains from fused regression are limited to those interactions which have fusion constraints, or if they extend to the entire network. To test this we used multiple 20 TF by 200 gene synthetic networks with varying proportions of orthologous TFs and genes. We divided networks into those interactions with fusion constraints (the constrained subnetwork) and interactions without fusion constraints (the non-constrained subnetwork). We varied the weight on the fusion penalty, lamS, and evaluated performance by computing AUPR on the constrained subnetwork, the non-constrained subnetwork, and the whole network (figure 1c).

Since the conserved subgraphs were similar to each other (more like figure 2a than 2b), we expected performance to improve as the fusion penalty weight increased. We observed this, particularly for the constrained subnetwork. As *λ*_*S*_ increased, interactions with fusion constraints were encouraged to be more similar. Interestingly, performance gains were seen even in the portion of the network that was unconstrained by fusion.

### 3.3 Adaptive fusion successfully identifies and unfuses ‘neofunctionalized’ genes

Orthology prediction is not a proxy for functional conservation [16, 55, 42]. To allow for orthologous genes to be unfused we implemented an adaptive fusion algorithm that attempts to optimize a nonconvex saturating penalty function on differences between fused interactions (figure 3). Pairs of interactions that are dissimilar even after fusion, which sit in the flat portion of this penalty function, are effectively “unfused,” and no further penalty is incurred as differences in interaction weights grow. Our network procedure strongly favors similarity of fused interactions, and only “unfuses” interactions when their similarity cannot be reconciled with expression data. As a result, the “unfusing” or relaxation of the fusion penalty on certain constraints is much more direct evidence for neofunctionalization than comparing separately fit networks could provide.

**Figure 3:**
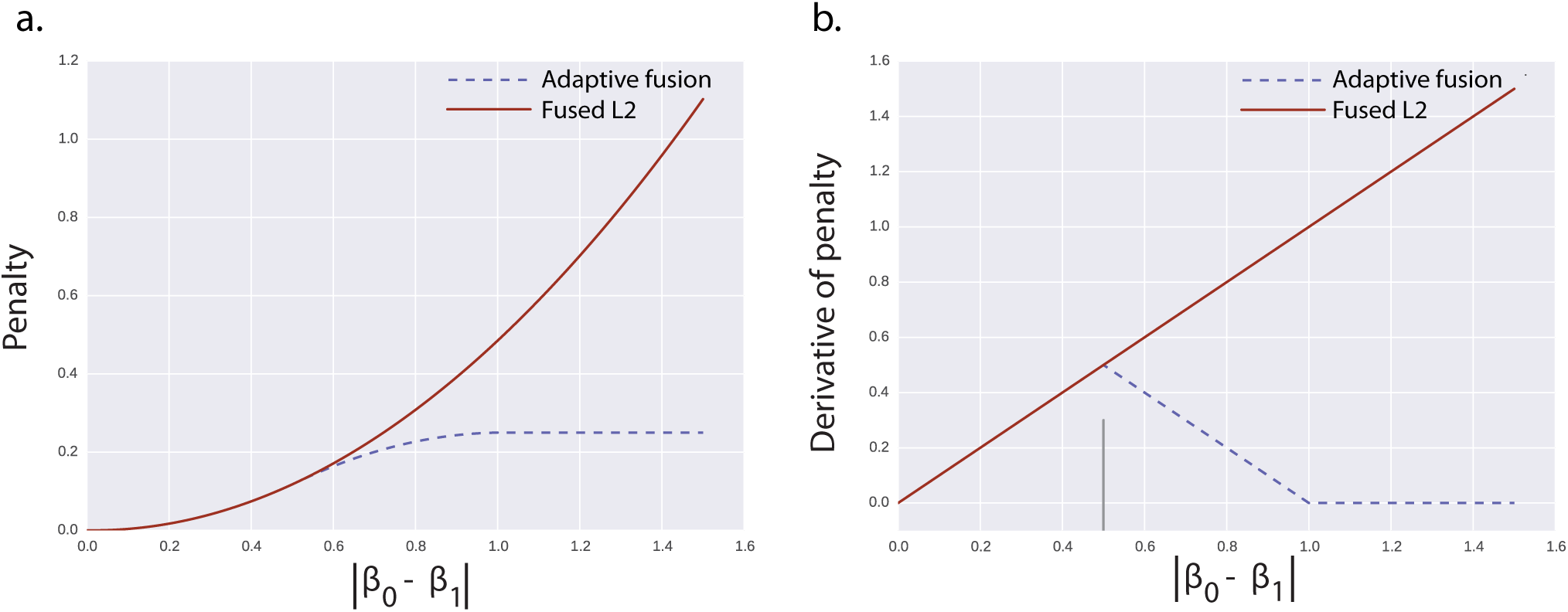
Adaptive fusion loss function (**A**) and derivative of loss function (**B**). **A**. Adaptive fusion is a quadratic around the origin, begins to taper at a/2, and plateus at a. After the plateu, increasing the difference in interaction weight of fused interactions does not further affect the penalty incurred through fusion. As a result, interaction weights in this zone are effectively unfused from one another (the fusion penalty behaves like a constant). **B**. Shows the derivative of the adaptive fusion penalty, which is used to implement adaptive fusion through local quadratic approximation. The adaptive fusion penalty is modified from SCAD (smoothly clipped absolute deviation) and MCP (minimax concave penalty) functions and like these penalties has a zero derivative far from the origin.

We performed a simulation to assess the ability of our adaptive fusion algorithm to distinguish which parts of two input networks are conserved vs. neofunctionalized (figure 4). We generated synthetic fused networks and introduced error in the fusion constraints by adding false positives and negatives to the orthology information given to the solver. Because we knew which entries in the orthology mapping were “incorrect” (not reflected in the generation of the networks), we could correctly label fusion constraints that involved one or more “incorrect” mappings. We verified that adaptive-fusion unfused mostly “incorrectly fused” interactions (figure 4a red dots), while leaving truly analogous interactions fused (figure 4a green dots).

**Figure 4:**
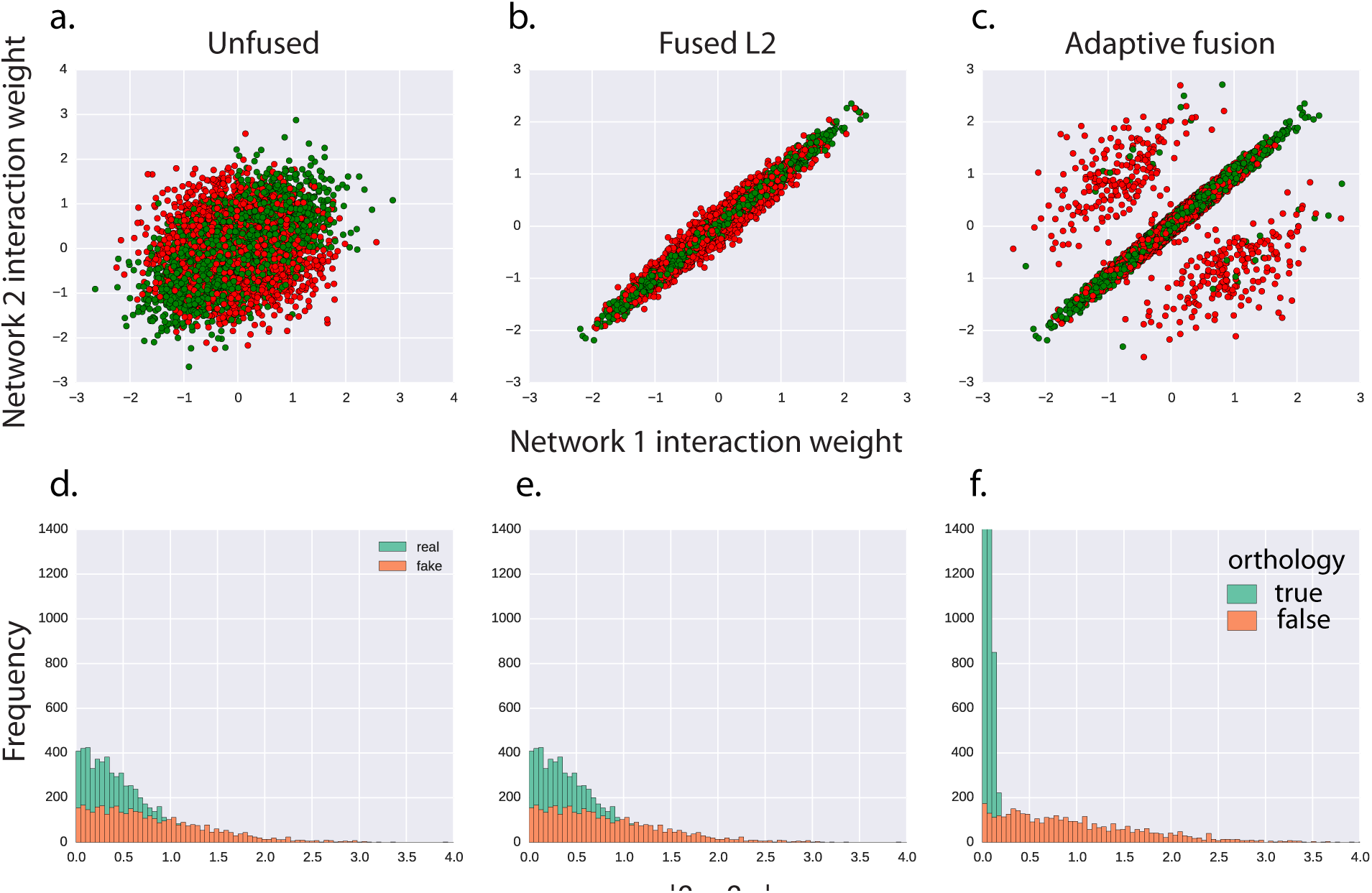
In order to evaluate the performance of adaptive fusion in unfusing interactions that have diverged through evolution, we performed a series of simulations inferring networks given a partially corrupted list of orthology mappings. Networks were generated with 35 TFs by 200 genes, 60% orthology coverage and 40% false orthology coverage. **A**. We plot the interaction weights between pairs of fused interactions in network 1 (x-axis) and network 2 (y-axis) following network inference without fusion (*λ*_*S*_ = 0). Interactions that are generated from false orthologs are marked as red, while interactions generated from true orthologs are shown in green. As expected, false fusion constraints give rise to uncorrelated weights (red dots). **B**. When fit with fused-L2, fusion constraints give rise to very similar weights in the two species for ‘true’ and ‘false’ interactions. **C**. Adaptive fusion run on the same network unfuses constraints for which the inferred weights are dissimilar beyond a certain point. Here, unfused interactions are almost entirely interactions between false orthologs. **D-F** Show the distribution of the absolute value of the difference in inferred weight for interactions with true fusion constraints (green) and false fusion constraints (orange)

We then compared the recovery of interaction weights which were accurately fused, and recovery of interaction weights which were inaccurately fused due to incorrect orthology information (figure 4b). Because fused L2 heavily penalizes large differences between weights which are predicted to be similar, it is able to retrieve a more accurate network for those interactions with true fusion constraints than by learning networks separately (measured by MSE between the true and inferred interaction weights). In this simulation, however, the gains accomplished through fused regression do not extend to those interactions lacking true fusion constraints, and the error remains similar to learning networks separately. When we applied adaptive fusion, we did not observe an improvement in network recovery (relative to fused L2). However, we were able to identify fusion constraints reflecting incorrect orthology information that had been provided to the algorithm (figure 4a).

### 3.4 Cross-species network inference using bacterial data

We used gene-expression data from *Bacillus subtilis* and *B. anthracis* in order to assess performance gains of fused regression on real data. Our *B subtilis* data set consists of 360 time-series and steady-state observations of 4891 genes, 4100 of which are protein coding [31], during the life cycle. Our B. anthracis dataset consists of 72 time-series and steady-state observations of 5536 genes comprising data from distinct points in the life cycle and iron-starvation conditions. There were 247 known transcription factors (TFs) in the *B. subtilis* dataset, and 248 TFs in the B. anthracis dataset. We obtained 1,870 one-to-one orthologs from Inparanoid [44], 95 of which are transcription factors, which produced 177,650 fusion-constraints between gene interactions within the two species. This number represents only 14.7% of the regulatory interaction matrix in *B. subtilis* and 12.9% in *B. anthracis*.

To assess network inference performance, and for use as priors, we used a gold standard of 3,040 known *B. subtilis* interactions with corresponding activation and repression sign. Of these 3,040 priors, 968 had corresponding interactions in *B. anthracis*. Based on our simulation results, we can expect the greatest gains in network-inference performance from fusion when the species of interest has a small number of available conditions, but data is abundant in a related species. However, in order to evaluate performance objectively a gold-standard of known interactions is necessary. As a result, we can only evaluate network recovery for *B. subtilis*, and *B. subtilis* also has the majority of our conditions. In order to simulate the data-poor regime, we subsampled our *B. subtilis* data. We divided our *B. subtilis* data into *k* folds, and then for each fold fit a network to the *B. subtilis* data from that fold alone fused to the entire 72 *B. anthracis* conditions (figure 5a). Though overall performance is hindered by our subsampling of *B. subtilis* data (a necessary procedure to allow evaluation of networks) we demonstrate marked improvement in learning the *B. subtilis* network when using fused regression (figure 5a). Notably, these performance gains occur mostly at low values of recall (near the top of our prediction ranks, where biologists would presumably focus validation and followup experiments).

**Figure 5:**
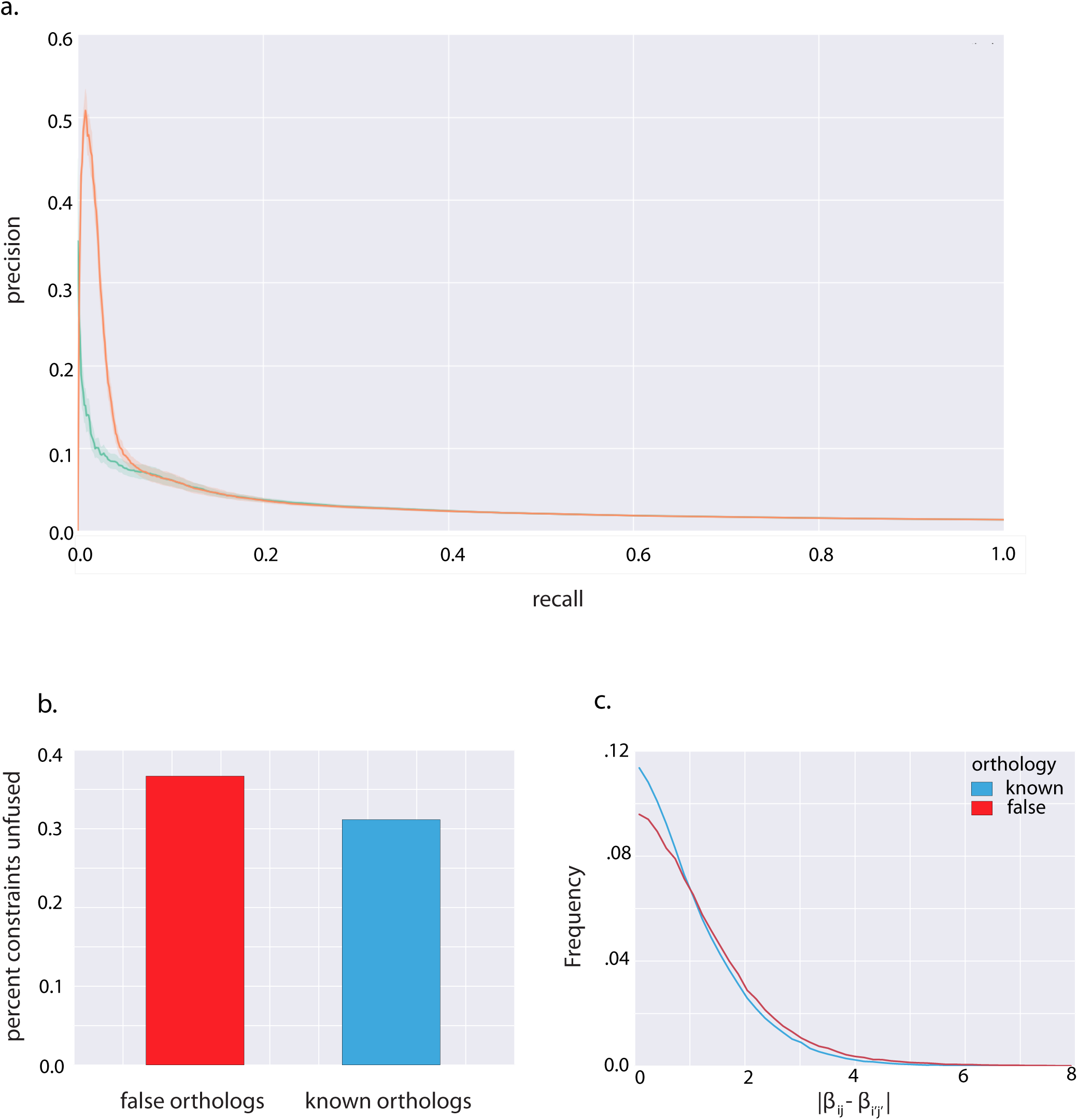
a. Using optimal lamR and lamP values, we use 20-fold cross validation to learn B. subtilis network. We comparedare performance when solving without fusion, and using L2 fusion with lamS = 1.0. Mean AUPR for lamS = 0 (unfused): 0.0298. Mean AUPR for lamS = 1: 0.0388. b. We test adaptive fusion using the same setup, with the addition of false orthology information. We set a 30% false potentialsitive rate, including 561 additional orthologs. We run adaptive fusion, setting the a term equal to the value above which 40% of constraints would unfuse, reflecting our belief that in addition to some of the known orthologs result in fusion constraints which should be relaxed. Here we show the percentage of constraints relaxed, from constraints created from the false orthologs and the known orthologs. We also show the distributions of differences in weights corresponding to fused constraints created from the false orthologs and the known orthologs, when networks are solved separately.

### 3.5 Testing adaptive fusion using bacterial data

The goal of adaptive fusion is to unfuse constraints between non-conserved interactions, while leaving intact all other constraints. However, because we lacked a comprehensive gold standard of known non-conserved interactions between *B. subtilis* and *B. anthracis*, we were unable to directly evaluate how accurately adaptive fusion identified these interactions. We opted instead to introduce a large number of random fusion constraints between genes not known to be orthologous. These fake constraints, which are unlikely to reflect any conserved network structure, served as a proxy for the unknown fraction of non-conserved interactions between orthologous genes. We ran adaptive fusion to learn networks for *B. subtilis* and *B. anthracis*, using these constraints, along with those generated by known orthology. We confirmed that our injected spurious fusion constraints were unfused at a higher rate than those generated by known orthologs (see figure 5b c). Although it may seem odd that a large fraction of fake constraints were left intact, we note that biological networks tend to be sparse, so that many of the random fusion constraints are between coefficients with near zero weight (and therefore near zero difference in weight) (figure 5c).

### 3.6 Integrating datasets from different platforms using fused regression

Although there are many large-scale collaborations which attempt to make protocols as uniform as possible for comparability between datasets generated by different labs [45, 30] and several methods for removing batch effects [23, 25], there still exists technical and biologial variability between many experiments attempting to capture the same or similar experimental conditions esspecially when experiments employ different experimental platforms. With the advent of RNAseq, for example, microarray based technologies are no longer the dominant assay for genome-wide expression, but a large body of accumulated legacy data remains useful if it can be integrated with more modern techniques. Currently, the most widely used approach to combining datasets for network inference is to learn networks from disparate datasets separately, then rank combine the networks as in Marbach et al [40]. We included, along with our *B. subtilis* dataset, a previously published dataset containing 269 samples covering 104 conditions, obtained using a different tiling microarray (vs custom microarray) and different strain of *B. subtilis* [43]. We compared performance when learning the networks separately and then rank combining (as in cite Ciofani) to learning the networks simultaneously using fusion regression and show large improvement in performance using our fused L2 approach (figure 6a).

**Figure 6:**
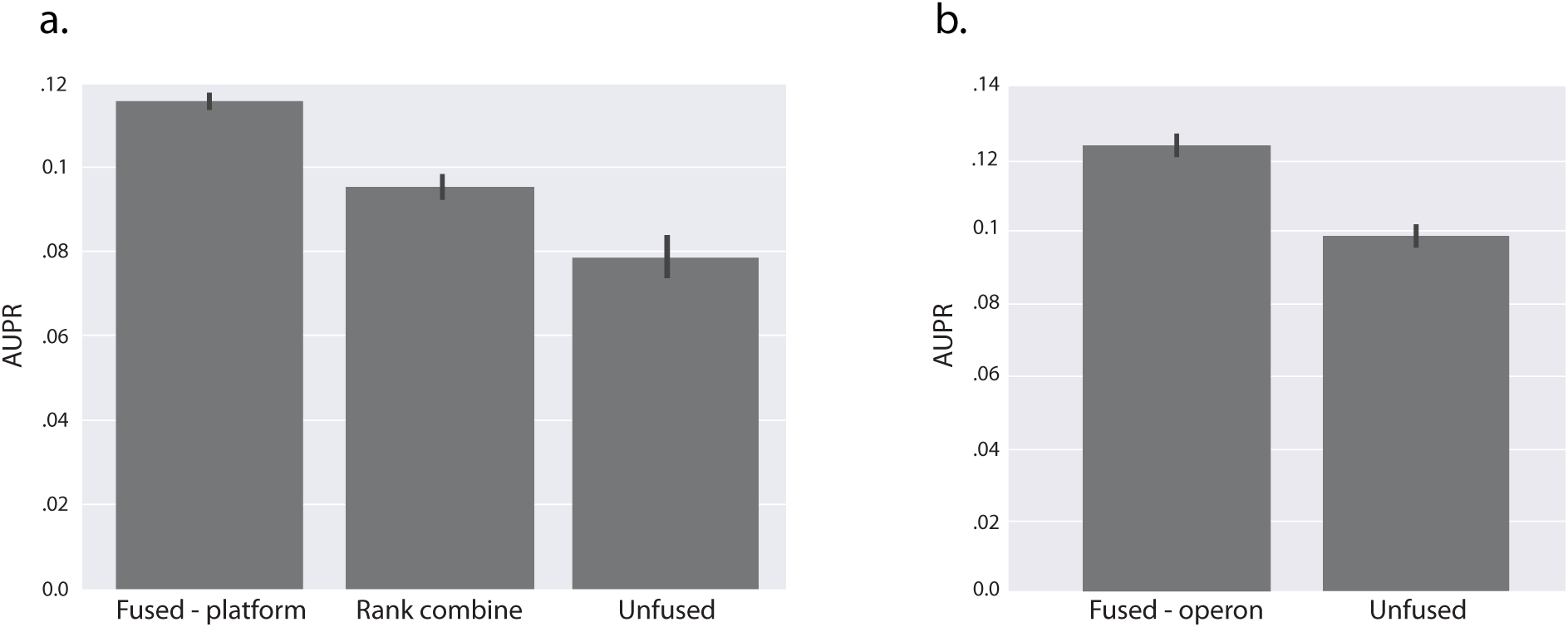
a. We use optimal lamR and lamP values, and solve for B. subtilis-1 and B. subtilis-2 networks using 10-fold cross validation. We fuse our original B. subtilis-1 dataset to another B. subtilis dataset, which we call B. subtilis-2, using orthology information, and evaluate performance of B. subtilis-1 using AUPR on gold standard. We compare L2 fusion with solving the networks separately without fusion, then rank combining as in Marbach et al., as well as solving B. subtilis on its own. b. We again use optimal lamR and lamP values, and solve for the B. subtilis network using 10-fold cross validation, and evaluate using AUPR on B. subtilis with gold standard. Here, we fuse genes in the same operon group and compare L2 fusion performance using operons with unfused network inference.

Information about the similarity of TF-gene interactions can also come from knowledge about the promoter region or the structure, for bacteria, of polycistronic transcripts. In bacteria, genes within the same operon are typically under the control of the same promoter [34]. We posited, therefore, that genes within the same operon will be regulated similarly by the same transcription factors. We applied fusion regression by creating fusion constraints between a given transcription factor and genes within the same operon, and showed a boost in *B. subtilis* network recovery using within-species fusion (figure 6b).

### 3.7 Transcription factor activity estimation integrates into fusion regression approach

We tested a combination of our fused regression approach with a method for estimating transcription factor activities (TFA). Rather than modeling gene expression using transcription factor mRNA abundance, we fit gene expression as a function of transcription factor activity, as applied to *B. subtilis* by Arrieta-Ortiz et al [1]. TFA activity estimates transcription factor activities that are modulated through mechanisms such as dimerization and interaction with required factors. TFA activity estimates have been shown prior to be better predictors of TF function than expression level alone in several contexts including similar network inference tasks[15] [1]. We estimate TFA based on known regulatory interactions using network component analysis [37]. To test the integration of this approach with our fused regression, we assessed the combination of *B. subtilis* datasets, as in figure 6a, with the incorporation of TFA estimation. We randomly divided the prior known interactions in half, and used half to learn TFA and to generate priors on network structure. The remaining interactions were reserved as a gold standard for validation. As in previous studies, we observed a marked improvement in network inference when using transcription factor activity (figure 7). We also obtained AUPR improvement when using fused regression on TFA, and showed that our gains from sharing information across datasets using fused regression were preserved and even enhanced by using TFA.

**Figure 7:**
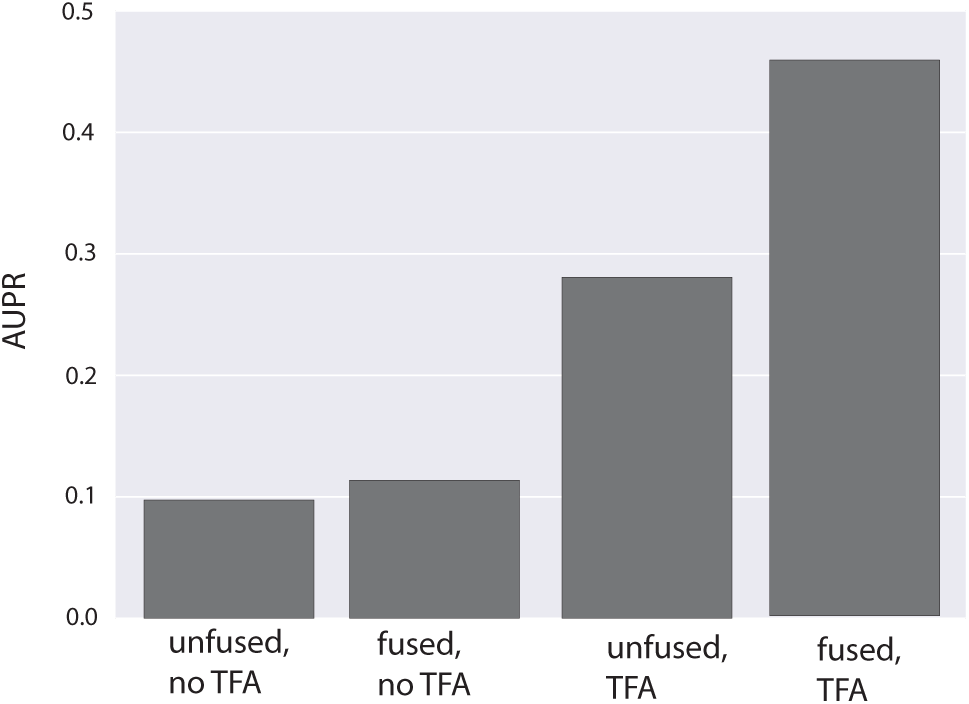
We integrate transcription factor activity (TFA) into our network inference, and solve for *B. subtilis*-1 and *B. subtilis*-2 networks using 10-fold cross validation, using fused L2 with and without TFA. As in figure 6, we evaluate performance on *B. subtilis*-1 using AUPR on gold standard.

## 4 Discussion

Gene expression data, such as microarray or RNA seq, provide information about the relationship between genes by allowing an experimenter to measure correlations in expression value over time or across conditions. Many sources of information - such as the knowledge that two genes are related through orthology or belong to the same operon - provide additional information about the relationships between these gene-gene relationships. For example, two genes that belong to the same operon are likely to have a similar set of regulators [34], but knowing that two genes are members of a polycistronic transcript does little to inform the identity (strength, sign) of those regulators. Meta-information about the structure of gene regulatory networks, specifically which pairs of interactions are *a priori* likely to be similar to one another, can provide a powerful set of constraints to improve network inference performance [50, 48]. We present a general framework for gene regulatory network inference that incorporates this meta-information - termed fusion constraints - and apply the technique to the problem of simultaneous inference of regulatory networks in multiple species (*B. subtilis* and *B. anthracis*).

We apply this algorithm to the problem of network inference in two distantly related biological organisms – *B. subtilis* and *B. anthracis* – and show that network recovery is improved through the introduction of fusion constraints between pairs of orthologous genes. Many previous methods for cross-species network inference operate on the conserved subset of orthologous genes [10]. This assumption may be appropriate with very closely related species, but could not be applied in this domain, where a large fraction (62% and 67%) of the *B. subtilis* and *B. anthracis* genomes do not have clear orthologs (and many orthologs have ambiguous many-to-many groupings). Our method, in contrast, can obtain improvements in network inference performance even when the conserved subset of genes is small. This approach is particularly interesting in light of the diversity of important model organisms used in modern biology. Different model systems provide different advantages and disadvantages for experimental design [54], but without a principled mechanism for combining data from multiple sources, it is difficult to fully leverage data obtained from even a slightly different model system. We further demonstrate the viability of fused L2 as a method for combining data from multiple experimental platforms, where fusion is between each identical regulator-gene pair. Because the algorithm we developed can accomodate constraints between arbitrary pairs of regulatory interactions, any biological prior representing information about expected regulatory similarity can be represented, even if the prior provides no information about the magnitude or direction of regulation. We demonstrate this flexibility through the novel incorporation of operon structure into the gene-regulatory network inference problem. In this application, fusion reflects the assumption that genes in the same operons have similar regulators [34]. The ability to incorporate multiple data sets describing related processes, as well as multiple data types, in a principled manner, helps us take advantage of the breadth of experimentation in biology to better learn the structure of gene regulation. We illustrate this by combining two different *B. subtilis* datasets and show that fused L2 is an improvement over current approaches to combining data [40] because of our ability to exploit the statistical power that our expanded datasets afford us.

Although it is important to take advantage of the similarities of related organisms for generating improved models of gene regulation, it is also critically important to understand how systems differ from one another. Our cross-species network inference method is premised on the assumption that orthologous genes have similar regulators. Existing approaches to the genome-wide testing of this assumption learn regulatory networks separately, then compare to identify conservation [2, 58]. Because network inference is typically underconstrained, fitting a network that describes a particular set of experimental observations involves sampling a single network from a large set of networks that fit the data equally (or almost equally) as well. As a result, the existence of a difference between corresponding regulatory interactions in a pair of experimentally derived networks is weak evidence that a difference truly does exist. Uncoupled global network inference algorithms are a very weak tool for uncovering evolutionary divergence. Our method explicitly favors recovering networks for which evolutionarily corresponding interactions are similar. As a result, the failure to obtain networks that confirm evolutionary conservation is stronger evidence that conservation does not exist; the next best network that does exhibit conservation must fit the data much worse to have overcome the bias built into the fusion constraints.

We have described a method – adaptive fusion – that attempts to learn which fusion constraints should be relaxed while the network is being learned. This method is based on minimizing a saturating penalty function on fusion constraints, similar to a class of penalties that have been developed to minimise bias in regularized regression [13, 62]. The result of adaptive fusion is both a network and a new set of fusion constraints, describing the learned fusion weights (including which fusion constraints have been relaxed). For the multiple species case, relaxation of fusion constraints represents orthologs which do not share similar interactions presumably due to evolution of regulatory circuitry [28]. When jointly learning networks describing processes in different cell lines, this may identify interesting context-specific behavior. Genes may be fused together on the basis of similar binding sites or chromatin features, and the relaxing of the fusion penalty indicates divergence of gene function.

Because our model shares its basic assumptions about the role of transcription factors in gene expression dynamics with models developed for single-species network inference, we are able to leverage techniques developed for the single-species estimation of transctiption factor activity [15]. The performance gains of this additional step in the cross-species case are significant. Our approaches – fused L2 and adaptive fusion – represent a very general framework for simultaneous network inference and the incorporation of structured biological priors. These priors – incorporated into our method as fusion constraints – allow the use of rich sources of biological knowledge, such as orthology and operon structure, which have informed experimental design, but are typically not incorporated into genome wide network inference algorithms. By accomodating the simultaneous inference of multiple related networks, we can improve network inference performance by allowing the efficient reuse of data from similar, but not necessarily identical, sources. A method for pooling data from multiple sources holds the promise of vastly expanding the quantity of data available for analysis, particularly in less commonly used model systems. At the same time these methods allow us to test our assumptions on how similar biological systems relate to one another, by allowing us to rule out conservation in a principled way, and at the genome-wide scale.

